# Genome-wide evidence for a hybrid origin of modern polar bears

**DOI:** 10.1101/047498

**Authors:** Tianying Lan, Jade Cheng, Aakrosh Ratan, Webb Miller, Stephan C Schuster, Sean Farley, Richard T Shideler, Thomas Mailund, Charlotte Lindqvist

## Abstract

Interspecific hybridization is recognized as a widespread phenomenon but measuring its extent, directionality, and adaptive importance in the evolution of species remain challenging. Polar bears possess unique adaptations to life on the Arctic sea ice, whereas their closest relatives -brown bears - are boreal and subarctic generalists. Despite largely non-overlapping modern distributions, genomic evidence demonstrates ancient admixture between these species. Here, we analyze new genomes from contemporary zones of species overlap as well as a previously sequenced 120,000-year old polar bear subfossil. We use explicit statistical fitting of data to admixture graphs to provide a framework for testing alternative scenarios of population relationships and gene flow directionality. Our analyses favor a single, parsimonious introgression event from relatives of extant Southeast Alaskan coastal brown bears into the ancestor of extant polar bears, which inverts the current paradigm of unidirectional gene flow from polar into brown bear. This conclusion has clear implications for our understanding of the impact of climate change: a specialist Arctic lineage may have been the recipient of generalist, boreal genetic variants at crucial times during critical phases of Northern Hemisphere glacial oscillations.

---

Recent research on ancient transfer of archaic human alleles into anatomically modern populations has highlighted important adaptive consequences of inter-species introgression. For example, ancient Denisovan allele states that introgressed into modern Tibetan human populations may confer altitude tolerance, and specialized lipid metabolic states in Europeans may derive from Neanderthal admixture^1,2^. Similar contact and admixture among other mammalian populations may have facilitated their own genetic adaptations in introgressed lineages, for example, during times of climate change. The high-Arctic polar bear (*Ursus maritimus*) and the lower-latitude brown bear (*U. arctos*) are recognized as highly distinct yet very closely related species^3,4^. However, recent research strongly points to ancient introgressive hybridization between these lineages^4-6^. Initially, this work centered on polar bear admixture with brown bears from Alaska’s Alexander Archipelago (the so-called ABC brown bears), since ABC brown bear mitochondrial haplotypes are closer related to haplotypes of polar bears than they are to mitochondrial haplotypes found in non-ABC brown bears^4,7-10^. Further analyses of nuclear genomes suggested widespread allele sharing among polar bears and various North American brown bears, albeit with closest proportion still found between polar bears and ABC brown bears^5,6^.

The current consensus scenario in the literature for brown and polar bear admixture involves multiple polar bear introgressions into brown bear lineages, possibly also including extinct Irish brown bears^11^. This hypothesis has implications for how climate change and range overlap may have influenced adaptive evolution by suggesting that generalist, boreal predators were the recipients of high-Arctic specialist alleles, with a selective barrier to gene flow likely to act in the opposite direction^5^. The converse, which has yet to receive empirical support, would implicate capture of boreal-adapted alleles by Arctic specialists known to be highly sensitive to climate change. Given these differences, the directionality of this admixture requires further scrutiny with a more complete sampling of crucial North American brown bear lineages and methodologies that permit explicit testing of alternative admixture hypotheses.

To evaluate alternative adaptive consequences of admixture polarity, we generated ten new bear genomes, including individuals from mainland Alaskan locales of contemporary brown and polar bear sympatry (Fig. 1 and Supplementary Information section I). Combining these new genomes with previously sequenced genomes of black (*U. americanus*), brown, and polar bears^4,6^ provided 25 genomes with up to 60-fold sequence coverage, representing all major brown bear maternal lineages. We aligned the sequence reads from these genomes to a polar bear genome assembly^12^ and called over 29.8 million nuclear single nucleotide polymorphism (SNP) genotypes that were filtered and prepared for downstream analyses (Supplementary Information section II). Phylogenetic analysis of assembled mitochondrial (mt) genomes (Fig. 2a) confirms previously reported findings of a close maternal relationship between polar and ABC brown bears, which in turn are sister to a brown bear from Finland (representing European clade 1^13,14^). Sister to this larger lineage are the remaining brown bear individuals from three main matrilines: a lineage comprising individuals from Yellowstone and Glacier National Parks (continental, or clade 4 bears^13,14^) and two sister lineages comprising Eurasian and Alaskan bears that represent Western and Eastern Beringian bears (or clades 3a and 3b, respectively^13,14^). The nuclear autosomal SNP phylogeny (Fig. 2b), as well as a phylogeny based on X chromosome SNPs (Supplementary Fig. 4), is incongruent with the mt phylogeny in that polar and brown bears comprise two distinct nuclear lineages. In the autosomal tree, all brown bears form a strongly supported clade, with a lineage of European brown bears (hereafter referred to as EBB) sister to a clade of Alaskan brown bears (BB bears), plus a monophyletic group of continental bears (YB) sister to the ABC brown bears. Interestingly, the Western and Eastern Beringian brown bear maternal lineages are not evident from the autosomal tree. The polar bear clade is poorly resolved internally, although the individuals from Svalbard group together as sister to a polar bear from Greenland. Principal component analysis confirms these major groupings (Supplementary Fig. 5 and 6).

**Figure 1.**
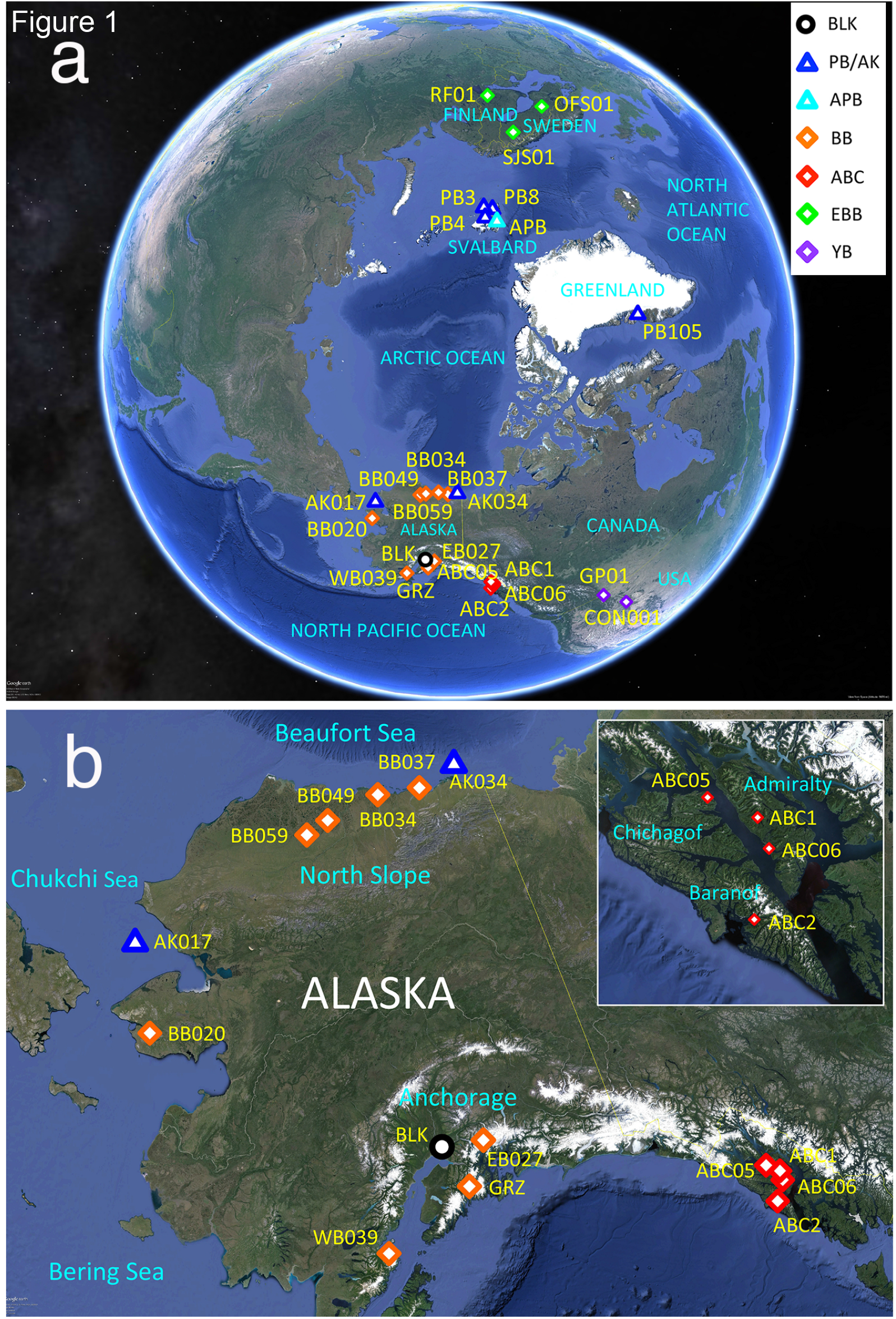
Maps showing localities of **a**, all samples included in this study and **b**, samples from Alaska only. The polar bears constitute three individuals from Svalbard (PB3, PB4, PB8), two individuals from Alaska (AK017, AK034), and one bear from Greenland (PB105). The brown bears comprise three European bears (SJS01, OFS01, RF01) and 14 bears from North America (BB020, BB034, BB037, BB049, BB059, EB027, WB039, GRZ, CON001, GP01, ABC1, ABC2, ABC05, and ABC06). One black bear from Alaska (BLK) was included in this study. See Supplementary Table 1 for more information.

**Figure 2.**
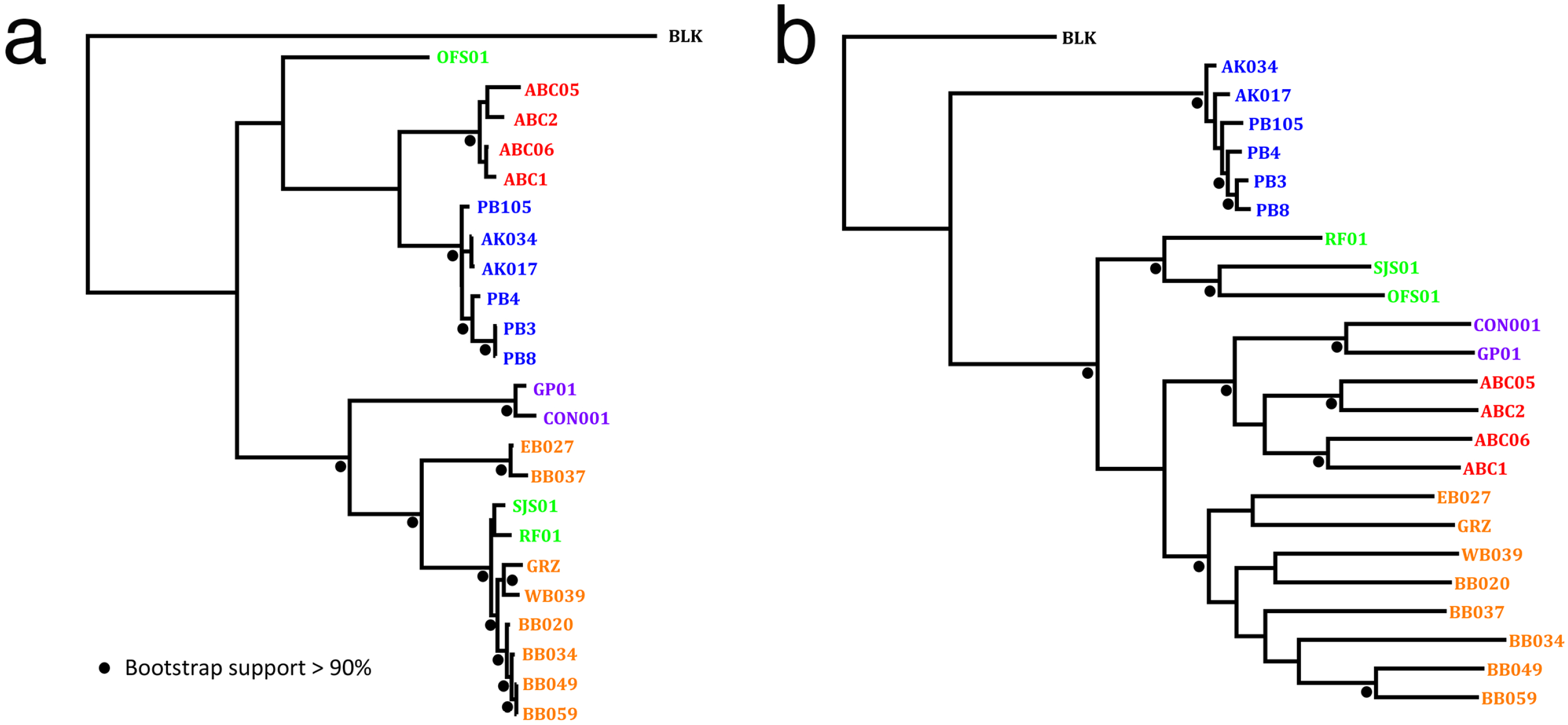
Phylogenetic maximum likelihood trees based on **a**, complete mitogenomes and **b**, autosomal SNPs.

Since we do not expect a simple, bifurcating tree to fully describe bear population interrelationships, we applied multiple measures to estimate genome-wide admixture^15-19^. We first produced a maximum likelihood drift tree using *TreeMix*^18^, which infers patterns of population splits and mixtures among multiple populations. Although the initial tree largely recapitulates the splits already seen (Fig. 2b), 0.45% of the variance is residual to the model’s fit, i.e., is not captured by the tree. Hence, we sequentially added five migration (admixture) events to the tree, and a number of potential admixture events stand out; these include interspecific exchange (1) between polar bears and the YB plus ABC brown bears, and (2) between YB plus ABC brown bears and an entity outside the brown/polar bear lineage that represents the edge leading to black bear (Supplementary Fig. 7). We also performed a cluster-based analysis of admixture^15^, and although we mostly observe admixture among the brown bear populations, at K=5 there is significant allele sharing among two polar bear individuals, a BB bear, and an EBB bear (Supplementary Fig. 8).

To obtain more detailed insight into admixture patterns and the population structure of the bears we employed the *f*_3_ and *f*_4_ statistics^19^, which use correlations in allele sharing to measure drift between populations or individual samples. The *f*_4_ statistic (closely related to the *D* statistic) is a four-taxon test of admixture that has become an important tool for estimating gene flow in population genomics^19-21^. This statistic measures the correlation in allele sharing among source and admixed target populations given an ancestral outgroup by computing the overlap of genetic drift paths in a phylogenetic tree that relates the populations^19^. The *f*_3_ statistic, or three-population test, is a formal test that can provide definitive evidence of admixture^19^. We calculated *f* statistics for all possible combinations of bear individuals and populations, also including genomic data from a 120,000-year old polar bear subfossil^5^ (Supplementary Information section III), to consider potential introgression between polar, brown and American black bear in all directions (Supplementary Information section IV).

When the *f*_3_ statistic is significantly negative it provides evidence that a population is admixed (although a positive value does not necessarily indicate non-admixture^19^). The only significantly negative *f*_3_ statistic was observed for the BB bears (Supplementary Fig. 16), where all comparisons, except those involving one individual (GRZ, from the Kenai Peninsula), show significantly negative values, particularly when juxtaposed against EBB and YB pairs. We conclude from these results that BB bears are admixed between (relatives of) the latter two populations. Using the *f*_4_ statistics and black bear (BLK) as outgroup, we evaluated levels of shared drift by comparing (1) each polar bear population against all pairs of brown bears, *f*_4_(BLK, polar; brown_1_, brown_2_) (Supplementary Fig. 17), and conversely, (2) each brown bear population against every pair of polar bear individuals, *f*_4_(BLK, brown; polar_1_, polar_2_) (Supplementary Fig. 18). In the first case, we did not observe any difference among choices of extant polar bears, whereas a striking gradient of polar bear likeness within the brown bears appeared. Brown bears from Baranof and Chichagof Islands (ABC-BC) exhibited the closest relationship with polar bears, followed in descending order by bears from Admiralty Island (ABC-A), then YB bears, which are slightly closer than BB bears, and finally EBB bears, which showed the least polar bear relatedness (Fig. 3a). This overall pattern is consistent with previous research that attributed the allele sharing to uneven amounts of polar bear admixture into different brown bear lineages^5^. When pairs of brown bear individuals are compared with the ancient polar bear (APB) instead of modern polar bear populations, however, we see less of a gradient and fewer significant values (Fig. 3a). In the second case, where each brown bear population is compared against every pair of polar bear individuals, we see no difference in *f*_4_ among choices of brown bear populations, with all extant polar bear pairs showing the same degree of shared allele frequencies with brown bears. The only sample that stands out is the ancient polar bear, APB, which shows less shared gene flow with brown bears than do extant polar bears (Supplementary Fig. 18). Such a pattern would be consistent with introgression of brown bear alleles into polar bear subsequent to the split between the ancient polar bear and the ancestors of the extant polar bears, and prior to the diversification of all extant polar bears. In this case, the observed gradient among brown bear populations described above would be interpreted as patterns of relatedness based solely on cladogenesis and drift.

**Figure 3.**
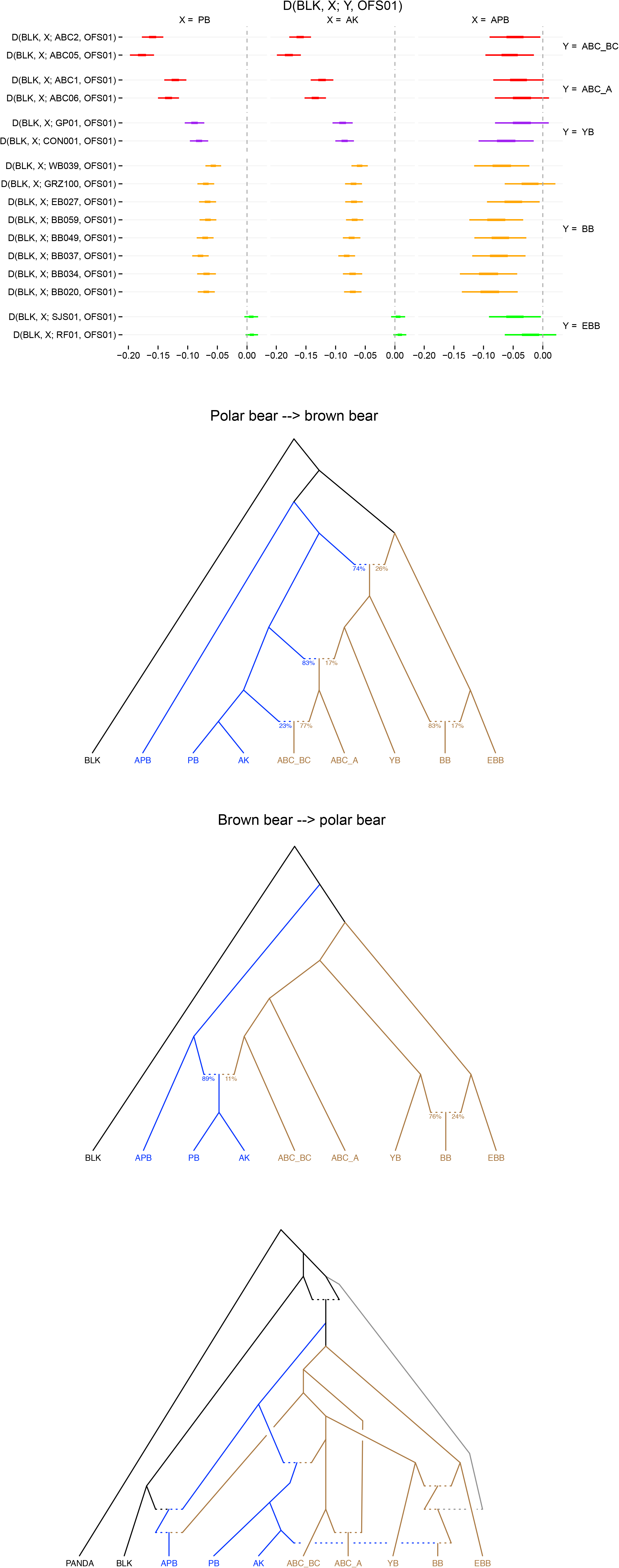
*f* statistics results and admixture graphs showing **a**, a gradient of allele sharing that is discerned when each polar bear population is compared against all pairs of brown bear populations, *f*_4_(BLK, polar; brown_1_, brown_2_) **b**, a scenario of population relatedness and gene flow, in which the observed gradient of shared drift in the *f*_4_ statistics are inferred as gene flow from polar bear into brown bear (left panel) and an alternative scenario, which receives a smaller sum of squared error, and where the observed *f*_4_ coefficients are inferred as gene flow from brown bear into polar bear (right panel). The admixture graph in **c**, is among the best fit graphs – based on the right graph in (**b**) - when shared drift with black bear is also considered.

Estimation of gene flow from black bear (BLK), using panda (*Ailuropoda melanoleuca*) as the outgroup, uncovered several clear patterns (Supplementary Fig. 19): the ancient polar bear, APB, received the most gene flow from BLK (or a relative of BLK), followed by YB, ABC-A, ABC-BC, extant polar bears, EBB, and finally BB bears. Within polar bears, PB bears from Svalbard appear to share more black bear alleles than AK bears from Alaska. Among the BB bears, two individuals stand out: WB039 (from Douglas River) appears to have less black bear gene flow than the other BB samples, and GRZ appears to have the most. Although as a population BB bears have the least gene flow from BLK, such patterns could indicate recent admixture events following BB divergence.

The *f* statistics by themselves do not predict the direction of gene flow. Admixture graph fitting, however, provides a rigorous test for whether a proposed evolutionary model fits the data^19,22^. Given such an admixture graph, the expected values of all *f* statistics can be expressed in terms of edge lengths (measured in drift) and admixture proportions. Conversely, given observed statistics and a graph topology, we can fit the edge lengths and admixture proportions to the data. We constructed admixture graphs and fitted them to values from our *f* statistics to model the necessary admixture events and the most likely evolutionary relationships (Supplementary Information section V). When considering the different *f*_4_ statistics as the result of introgression from polar bear into brown bears, we can fit this to an admixture graph only by invoking at least three admixture events (Fig. 3b and Supplementary Fig. 23) and with a sum of squared error (SSE) of 0.209. Surprisingly, if we conversely interpret the *f*_4_ statistics as the consequence of gene flow from brown bears into polar bears, which conflicts with most recent interpretations^5-7^, we can fit the observed statistics with smaller error (SSE=0.029) and more parsimoniously with fewer admixture events (Fig. 3b and Supplementary Fig. 24). This strongly suggests that the better fitting model involves gene flow from brown bears into polar bears rather than in the opposite direction. The favored graph also includes results from the *f*_3_ statistics that suggest BB is likely admixed between YB and EBB, and it posits that the ABC-BC population contributed more gene flow to the modern polar bear stem lineage than the ABC-A population did. However, other scenarios for the ABC brown bears with similar fits can be invoked, including a model in which an ancestral ABC lineage contributed gene flow to modern polar bears, and ABC-A bears later received gene flow from another brown bear lineage, pulling them farther away from polar bears compared to the ABC-BC bears (Supplementary Fig. 26). This latter scenario receives some support from evidence that the ABC brown bear maternal haplogroup was formerly more widespread^11,23^, likely even trans-Beringian (Lan and Lindqvist, unpublished data).

Next, we fitted possible black bear gene flow onto our best-fit admixture graph by adding panda as outgroup (Supplementary Figs. 30 and 31). Such admixture has precedence in the literature, since cytonuclear incongruence in the phylogenetic placement of Asian (*U. thibetanus*) and American black bears has previously been observed^24^, as have high levels of shared polymorphisms between brown and black bears^25^. Given a strong indication that APB shares alleles with BLK, we first added an admixture event for that relationship (Supplementary Fig. 30). The remaining statistics not predicted correctly by the graph mostly involve cases where BB bears are less related to BLK than the other bears are (Supplementary Fig. 30c). Rather than invoking individual gene flow events from BLK into each non-BB population, we can achieve such a pattern by postulating that a ghost population from outside the brown, polar, and black bear clade introgressed into the BB lineage, hence diluting any BLK gene flow (Supplementary Fig. 31). We also tested an alternative scenario involving a ghost population that branches off the ancestor of all brown bears, followed by ancient admixture between black and brown bear (Supplementary Fig. 33). In that case, introgression from the ghost lineage into BB would give the same pattern of *f*_4_ statistics. Although this alternative did not improve the fit of the graph, it is still among the best fitting graphs. Such admixture events with ghost populations could reflect hybridization with known species that went extinct at the end of the Last Glacial Maximum, such as the giant short-faced bear (*Arctodus simus*) in the first scenario, or the Etruscan bear (*U. etruscus*) or European cave bear (*U. spelaeus*) in the alternative.

Since fitting the data perfectly would require us to split all populations into single individuals, rendering graph fitting unfeasible, we examined alternative population-level graphs by modeling different relationships among established populations. Among 37 alternative graphs (Supplementary Fig. 32), five had the smallest error at the population level (0.0164), although several graphs had only slightly greater errors (Supplementary Table 7). Although it is not possible to test the significance of fits on these alternative graphs, their general models of population relationships are very similar. The models mainly differ in admixture proportions and the sequence of admixture events for the BB population (which includes three waves of gene flow that do not spill over to other populations), different scenarios for the ABC bears (see above), and whether or not brown bear gene flow into APB is ancestral to all modern polar bears. The latter (Fig. 3c) would be consistent with the observation that all polar bears have mt genomes that branched off from brown bear very recently (Fig. 1a)^8^. The estimated admixture proportion between the ABC brown bears and the ancestor of modern polar bears (11%; Fig. 3b) is in accordance with previous estimates^4,5^. Gene flow from EBB bears and signals of differential admixture within BB bears might reflect a dynamic biogeographic history with several migration waves of brown bears into the New World^13^, as well as more recent gene flow with polar bear. Nevertheless, all BB bears, as well as the EBB and ABC bears, exhibit largely similar fluctuations in historical effective population size, which stands in sharp contrast to the patterns of effective population size history of either polar bears or black bear (Supplementary Fig. 9).

Many organisms show shifts in latitudinal range in response to changing global climate^26^, and colonizing species have in some cases been shown to capture local adaption by hybridizing with closely-related resident lineages^27^. As a result, hybridization may catalyze adaptive evolutionary change^28^. Hybridization requires sympatric distribution during the breeding season. Although contemporary ranges of polar bears and their lower-latitude closest relatives are discrete across much of the Arctic, sympatric distribution during the breeding season occurs along the Arctic coast west of Hudson Bay to the northern Chukchi Sea coast. Observations over the last decade indicate that brown and black bears are moving northward into the Canadian Arctic Archipelago^28^. Furthermore, polar bears are increasingly summering in nearshore terrestrial and barrier island habitats in the central Beaufort Sea, perhaps facilitated by the presence of fall bowhead remains^29,30^ but more likely due to the loss of nearshore Beaufort Sea ice during summer^29-32^.

Results of our admixture graph fitting, which provide a powerful framework to test alternative scenarios of population interrelationships, particularly when critical samples are included, support an inverted paradigm shift: past introgression from brown bear into the ancestor of modern day polar bears. This not only reconciles the highly paraphyletic nature of brown bear maternal lineages^8,11,23^, but also has strong relevance for our understanding of potential adaptive responses to climate change. Our data suggest that following the divergence between ancestors of black, brown, and polar bears, introgression events among these species involved significant gene flow *into* the Arctic lineage at least twice from ancestors of extant brown bears and once from ancestors of extant black bears, possibly facilitating the capture of novel genes by Arctic specialists (polar bears) from colonizing boreal generalists (brown and black bears). Although there is likely strong purifying selective pressure on polar bear phenotypic features adapted to extreme Arctic life, novel, heritable traits transferred to polar bears could have become selectively advantageous during certain past periods of climatic change. Insight into the potential adaptive importance of ancient polar-brown bear admixture could come from regions where modern polar bear genomes are more similar to brown bears than they are to the ancient polar bear predating this admixture event. The collection of more complete ancient DNA data than we present here, however, will be required to infer any potential genome-wide adaptive signals.

Despite a predominant pattern of gene flow from brown bear into ancestors of modern polar bears, we also show evidence that mainland Alaskan brown bears (BB) may be secondarily admixed with polar bear alleles, and indeed, it seems likely that the true history of gene flow between these species has been multidirectional. We hypothesize that the direction of introgression may have been correlated with climate oscillations. Gene flow may have been predominantly from boreal populations (brown bears) into Arctic populations (polar bear) during warming periods, with Arctic specialists in part adapting by capturing generalist alleles. Conversely, during cooler periods when polar bear range expanded, introgression may have occurred from the Arctic lineage into the boreal species, thereby facilitating adaptation of the latter through capture of Arctic-adapted alleles. Genetic patterns from such scenarios are clearly obscured by a complex brown bear biogeographic history, and our results indeed suggest distinct, lineage-specific geographic expansions of brown bears into the New World, i.e., BB independently from YB plus ABC (Fig. 4). Only more complete genomes, including from ancient bear remains, will permit proper modeling and timing of these admixture events and demographic expansions, which in turn will better inform their correlation with events during Earth history.

**Figure 4.**
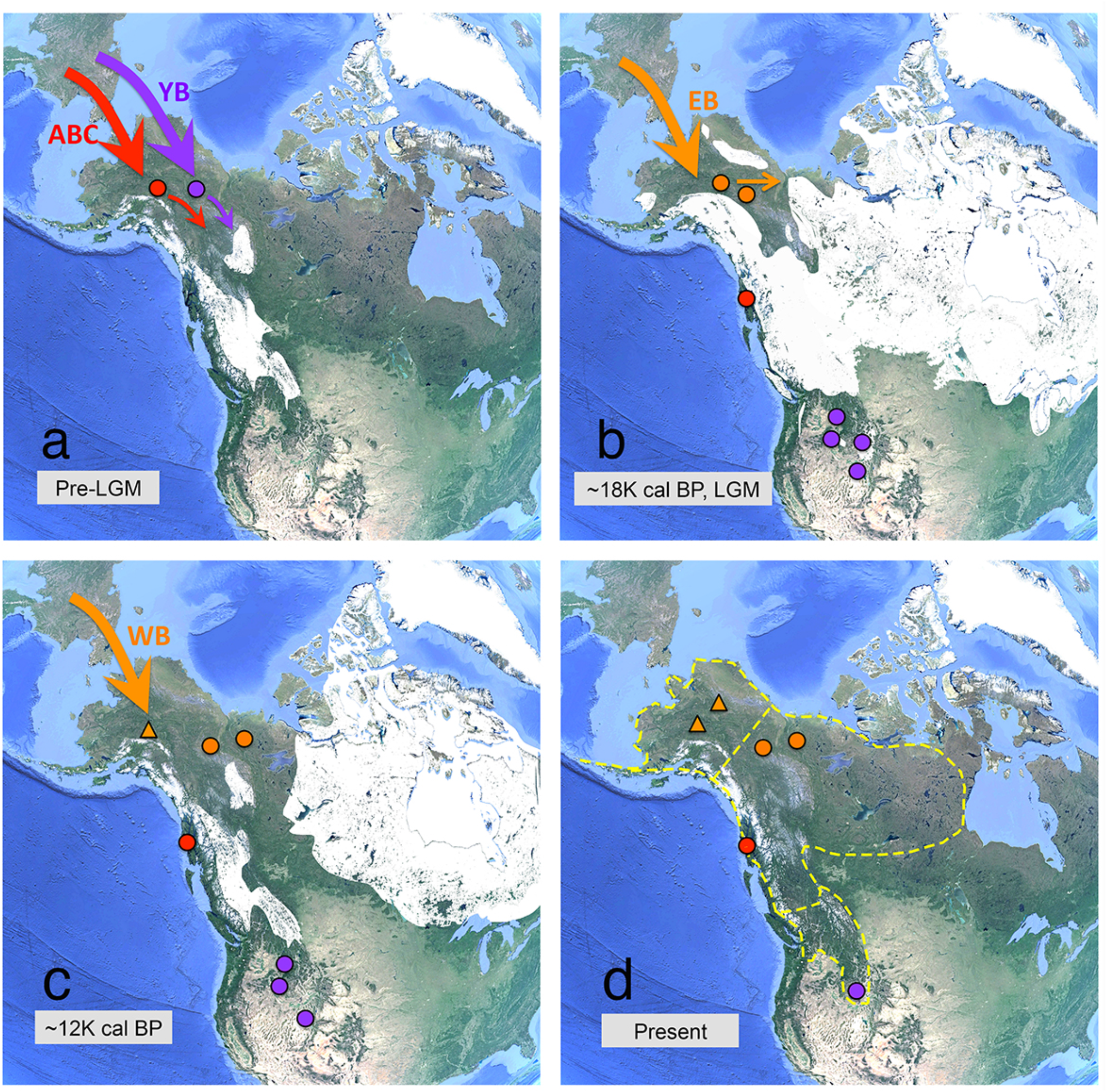
Model of biogeographic scenarios of brown bear movements into the New World based on analyses of nuclear and mitochondrial genomic data. **a**, pre-LGM (Last Glacial Maximum); **b**, circa 18K cal BP; **c**. circa 12K cal BP; d. present. Brown bear populations are color-coded as: ABC: red circle; YB: purple circle; EB (Eastern Beringian): orange circle; WB (Western Beringian): orange triangle. The extent of glaciation at circa 18K (**b**) and 12K cal BP (**c**) is reproduced from Dixon *et al*.^43^, and the distribution of modern brown bear clades (dashed lines in panel **d**) is reproduced from Waits *et al*.^10^ and Davison *et al*.^13^.

## Methods

Detailed sampling and experimental procedures can be found in Supplementary Information.

### Sampling, DNA Extraction and Genome Sequencing

For sequencing of new bear genomes, blood and tissue samples were collected from one brown bear from Yellowstone National Park, seven brown bears from throughout Alaska, and two polar bears from Chukchi and Southern Beaufort Sea in Alaska (Supplementary Table 1 and 2). All samples were collected from wild populations using standard procedures. DNAs were extracted using the protocols reported by Medrano *et al*.^33^ with modifications reported in Sonsthagen *et al*.^34^, or using the DNeasy Blood and Tissue Kit (QIAGEN, USA) according to the manufacturer’s specifications. DNA libraries were prepared by using the Illumina TruSeq Nano DNA Library kit and then sequenced to generate 150bp paired-end reads by using the HiSeq2500 Rapid Sequencing platform at the Singapore Centre on Environmental Life Sciences Engineering Sequencing statistics are provided in Supplementary Table 1.

### Mapping, SNP Calling and Filtering

Illumina reads from previously sequenced bear samples^4,6^ were downloaded from NCBI Sequence Read Archive (SRA) database, including one ancient polar bear, four modern polar bears, nine brown bears and one black bear. All paired-end reads from both the newly sequenced and previously sequenced bear samples were trimmed with Trimmomatic^35^ to remove adapters and low quality bases. Trimmed reads were then aligned to the draft polar bear nuclear genome^12^ and a brown bear mitochondrial genome (GenBank accession AF303110.1) using Burrows– Wheeler alignment (BWA)^36^ with default parameters. PCR duplicates were removed using MarkDuplicates tool from Picard suite (http://broadinstitute.github.io/picard/). Mapping statistics are provided in Supplementary Table 1. Nuclear genome and mitochondrial genome consensus sequences were called using mpileup in SAMtools^37^ with the option “-C 50”. SNP calling were conducted by using HaplotypeCaller in GATK toolkit^38^ with default settings. SNPs were filtered by depth, genotype quality, minor allele frequency, scaffold location and scaffold size using VCFtools^39^. Detailed filtering criteria and datasets used for analyses are described in Supplementary Information section I.

### Analytical Procedures

Phylogenetic analyses based on complete mitochondrial genomes, autosome-associated SNPs dataset, and X chromosome-associated SNPs dataset were conducted using RAxML^40^ with 1000 bootstrap replicates. Principle component analyses were performed using smartpca included in the Eigensoft package^41^ to infer the population structure and relationships among the bear samples. *TreeMix*^18^ analyses were performed allowing up to five migration edges on the maximum likelihood tree to infer the relationships, divergence, and major mixtures between populations. ADMIXTURE analyses with different K values ranging from 2 to 7 in an unsupervised mode were conducted for estimating individual ancestries and genetic structure. To analyze bear demographic history, pairwise sequentially Markovian coalescent (PSMC)^42^ analyses were implemented for all modern bear samples. See Supplementary Information Section II for more details.

We analyzed population structure of the bears using ADMIXTOOLS^19^ and *f* statistics, which uses correlations in allele sharing to measure drift between populations or individual samples. For the *f*_3_ statistics, *f*_3_(C;A,B), we only consider cases where A, B, and C come from different populations. The *f*_3_ statistics can capture drift on overlapping paths from C to A and C to B, and if C is admixed, some of this drift can contribute a negative value to the statistics. We carried out the *f*_4_ statistics, *f*_4_(W,X;Y,Z), to test for admixture between different populations. The *f*_4_ statistics, where we assume a topology (W,(X,(Y,Z))), is negative if there has been (more) gene flow between X and Y than between X and Z (or more gene flow between W and Z than between W and Y) since the split between Y and Z, and positive if there has been more gene flow between X and Z than between X and Y. Admixture graphs, which fit the edge lengths and admixture proportions to the *f*_4_ statistics outcomes, were constructed by using the admixturegraph R package (https://github.com/mailund/admixture_graph) to visualize the genetic drift and gene flow among our bear populations. Also see Supplementary Information sections IV and V for more details.

## Acknowledgments

We thank G. K. Sage and Daniela Drautz-Moses for laboratory assistance. This work was supported by the National Fish and Wildlife Foundation (to C.L.) and Alaska Department of Fish and Game (to S.F. and R.T.S.).

